# Long-term ovarian cancer survivors: spatial transcriptomics depict ligand-receptor crosstalk heterogeneity at the tumor-stroma interface

**DOI:** 10.1101/2022.06.04.494814

**Authors:** Sammy Ferri-Borgogno, Ying Zhu, Jianting Sheng, Jared K. Burks, Javier Gomez, Kwong Kwok Wong, Stephen T.C. Wong, Samuel C. Mok

## Abstract

Advanced high-grade serous ovarian cancer (HGSC) is an aggressive disease that accounts for 70% of all ovarian cancer deaths. Nevertheless, 15% of patients diagnosed with advanced HGSC survive more than 10 years. The identification of predictive markers associated with tumors developed from these long-term survivors (LTS) is crucial to identifying therapeutic targets for the disease, and thus improving patient survival rates. Reports to date have not fully established the stromal heterogeneity of the tumor microenvironment (TME) in ovarian cancer and its association with clinical outcomes. We used a spatial transcriptomics platform to generate spatially resolved transcript profiles in treatment naïve advanced HGSC from LTS and short-term survivors (STS), and determined whether cancer-associated fibroblasts (CAFs) heterogeneity is associated with survival in patients with advanced HGSC. We integrated spatial transcriptomics with single-cell RNA sequencing data to distinguish tumor and stroma regions, and developed a method to investigate spatially resolved ligand-receptor interactions between various tumor and CAF subtypes in the TME. In addition, we used multiplex immunohistochemistry techniques to validate our findings. We found that a specific subtype of CAFs and its spatial location relative to a particular ovarian cancer cell subtype in the TME correlate with long-term survival in advanced HGSC patients. We also demonstrated that significant APOE-LRP5 crosstalk occurred at the stroma-tumor interface in tumor tissues from STS compared to LTS, suggesting that such crosstalk plays a crucial role in modulating the malignant phenotype of HGSC, and could serve as a predictive biomarker of patient survival.

## INTRODUCTION

Advanced high-grade serous ovarian cancer (HGSC) is the most common ovarian cancer histotype and accounts for more than 70% of ovarian cancer deaths making it the most lethal gynecologic malignancy. HGSC is predicted to cause 12,800 deaths in 2022 in the United States alone (*1*). HGSC typically presents as aggressive advanced-stage disease (*2*) and is initially sensitive to platinum-and-taxane–based chemotherapy (with a 75% response rate) delivered before or after debulking surgery. However, the vast majority of patients with advanced HGSC (>75-80%) have recurrence 12 to 24 months after initial treatment, experience rapid disease progression, and die of progressively chemotherapy-resistant disease (*3, 4*). Nevertheless, 15% of patients diagnosed with advanced HGSC have overall survival (OS) durations of more than 10 years (*5, 6*).

Reports to date have not fully established the heterogeneity of the tumor microenvironment (TME) in ovarian cancer and its association with clinical outcomes. The TME, which is composed primarily of fibroblasts, endothelial cells, lymphocytic infiltrates, and extracellular matrix proteins, can directly affect cancer cell growth, migration, and differentiation and thus presents a unique opportunity for cancer diagnosis and treatment (*7*). The immune system is an important determinant of the TME; ongoing inflammation results in various immunologic gene products that create a favorable microenvironment for tumor growth and progression (*8–12*), and the presence of specific immune cell types, such as intratumor CD8^+^ T cells, is associated with improved survival in patients with various types of cancer, including ovarian cancer (*8, 9, 13*). These findings suggest that immune cell heterogeneity plays an important role in conferring the malignant phenotypes of cancer cells. In addition to immune cell heterogeneity, cancer-associated fibroblast (CAF) heterogeneity may also play essential roles in modulating tumor growth. CAFs are characterized by their expression of traditional markers, including alpha-smooth muscle actin (αSMA), S100A4, vimentin (VIM), fibroblast activation protein (FAP), and platelet-derived growth factor receptor alpha (PDGFRα) and beta (PDGFRβ) (*14*). Differential expression of these CAF markers has recently been found in the TME (*15*). Furthermore, TME has been found to contain additional CAF subtypes that have distinct biomarker expression profiles and cellular functions that differ from those of CAFs outside the TME (*16, 17*). However, the molecular mechanisms underlying the promotion or inhibition of cancer by various CAF subtypes as well as the interplay between different CAF populations and their spatial locations within the TME in ovarian cancer are not fully understood.

Recent innovations in high-throughput analyses of patient-derived specimens may address the clinical challenges described above. The application of integrated “omics” techniques (including proteomics, transcriptomics, metabolomics, and immunomics) has improved our understanding of how genetic changes affect the gene expression profiles (*18–20*). However, because most of the data generated with these techniques are derived from bulk tumor tissue, they have limited utility in the clinical management of HGSC. Previous studies have shown that the stroma admixture affects the interpretation and reproduction of molecular subtypes and gene signatures derived from bulk tissue (*21*). Recently, researchers used single-cell RNA-seq (scRNA-seq) analyses to characterize the heterogeneity of the HGSC TME, thereby providing valuable information about different HGSC subtypes and potential novel therapeutic approaches (*22*–*24*). However, the aforementioned techniques and analyses cannot provide spatial information. Increasing evidence in multiple cancer types suggests that the spatial location of various cellular components of the TME and their position in relation to tumor cells, immune cells, and blood vessels can modulate anti- and pro-tumor responses (*25*–*28*).

Spatial transcriptomics technology, which captures genome-wide readouts across biological tissue space, enables researchers to determine how genes are spatially expressed in the complex TME (*29*). In this study, we used spatial transcriptomics to demonstrate the prognostic significance of CAF heterogeneity in HGSC. We integrated spatial transcriptomics with scRNA-seq to distinguish tumor and stroma regions. In addition, we developed a method to investigate region-specific ligand-receptor interactions between HGSC and neighboring CAF subregions and subsequently identified a ligand-receptor pair in a subtype of CAFs and their neighboring HGSC cells that has prognostic significance.

## MATERIALS AND METHODS

### Patient samples

A total of 4 frozen and 27 paraffin-embedded tumor tissue samples obtained from patients with advanced (stage IIIB-IV) HGSC were used in the spatial transcriptomics and multiplexed immunofluorescence (mIF) experiments, respectively. Tissue samples were obtained from the ovarian cancer repositories at The University of Texas MD Anderson Cancer Center. They were collected from previously untreated patients undergoing primary cytoreductive surgery for ovarian cancer. After surgery, patients received platinum-based combination chemotherapy. Optimal surgical cytoreduction was defined as a residual tumor of no more than 1 cm in diameter. The overall survival duration was measured from the date of diagnosis to the date of death or censored at the date of the last follow-up examination. Short-term survivors (STSs; samples A10 and A12) were those with an overall survival duration of less than 24 months, and long-term survivors (LTSs; samples A4 and A5) were those with an overall survival duration of more than 120 months. Clinical data, including age, cytoreduction status (optimal vs. suboptimal), and overall survival, were obtained from the patients’ medical records. All samples and clinical data were collected with the approval of MD Anderson’s Institutional Review Board.

### Spatial transcriptomics

Spatial transcriptomics experiments were performed according to the manufacturer’s protocol (10x Genomics). Briefly, a 10-μm-thick HGSC tissue frozen section was placed onto a spatial transcriptomics expression slide to fit an 8 x 8 mm spatially barcoded array with 1007 spots, each with a diameter of 100 μm and a center-to-center distance of 200 μm. The spatial transcriptomics slide was then stained with hematoxylin & eosin (H&E) and imaged with the EVOS M7000 Imaging system (Thermo Fisher Scientific). Tissue permeabilization and cDNA synthesis were then performed directly on the tissue section. The pooled barcoded and spatially transcribed cDNA libraries for each sample were sequenced using Illumina NextSeq500 flow cells at MD Anderson’s Advanced Technology Genomics Core. Raw output base call (BCL) files from the sequencer were demultiplexed into fastq files (10x Genomics).

### scRNA-seq analysis and spatial transcriptomics RNA-seq analysis

See Supplementary Data for details on scRNA-seq and Spatial transcriptomics RNA-seq analysis.

### CAF annotation

The CAF clusters were annotated if the stroma clusters significantly expressed the traditional CAF markers (p < 0.01) (Figure 3A). For investigating the CAF heterogeneity across samples, the average log fold-change of the CAF genes in each CAF cluster was computed as the log fold-change of the average gene expression in that CAF cluster compared to the average gene expression in all other clusters in that sample.

### Multimodal intersection analysis

With the gene sets extracted from spatial transcriptomics and scRNA-seq modalities, the overlap between the region-specific (differentially expressed genes as identified in spatial transcriptomics data analysis above) and cell type specific gene sets (differentially expressed genes as identified in scRNA-seq analysis above) was then computed. We performed a hypergeometric test to identify the significant enrichment (using a threshold of p < 1 × 10^-10^) of any specific cell type within a tumor/stroma subregion (*31*) using 16,522 background genes.

### Gene Ontology overrepresentation analysis

Gene Ontology (GO) terms were downloaded from DAVID (Database for Annotation, Visualization and Integrated Discovery). GO terms for biological processes at 2-4 levels of granularity were used. The GO terms with less than 10 genes were removed. Details of the GO analysis are provided in the Supplementary Data.

### Ligand receptor interaction analysis

Using databases from Choi et al. and Yeung et al. (*32, 33*), which included 2,671 ligand-receptor pairs, we analyzed ligand-receptor interactions at the interface between every stroma and tumor subregion. At each interface, only the stroma spots and tumor spots that were nearest to the interface were considered. (The minimum distance of each stroma spot to the tumor and the minimum distance of each tumor spot to the stroma were equal to the minimum distance between the ST spots; that is, 200 μm.) Details of the ligand receptor interaction analysis are provided in the Supplementary Data.

### Multiplexed immunofluorescence

See Supplementary Data for details on multiplexed immunofluorescence experiments and analysis.

### Statistical analysis

We used a non-parametric Wilcoxon rank sum test to detect differences in the mean intensities/stroma area ratios and positive cells/total cells ratios between LTS and STS samples in the mIF experiments and to identify the differentially expressed genes in the spatial transcriptomics and scRNA-seq data analyses. A p-value less than 0.05 was considered statistically significant. For the analyses of differentially expressed genes of ST and scRNA-seq, adjusted p-value was computed based on bonferroni correction using all genes in the dataset.

### Data availability

All data needed to evaluate the conclusions in the paper are present in the paper and/or the Supplementary Materials. The scRNA-seq dataset was downloaded from GEO database under accession GSE118828 (*34*).

## RESULTS

### Generation of spatially resolved RNA-seq profiles of HGSC tumors

To generate spatially resolved transcriptomes from various cellular compartments of the ovarian TME, we performed spatial transcriptomics analysis on four samples of treatment-naïve advanced HGSC (Fig. 1A). We first mounted the tissue cryosections onto the spatially barcoded spatial transcriptomics microarray slides and performed H&E staining and brightfield imaging. The samples were then processed for spatial transcriptomics analysis; this processing included complementary DNA (cDNA) synthesis, amplification by *in vitro* transcription, library construction, and sequencing. We demultiplexed the sequenced reads and identified their spatial location within the tissue using the spatial transcriptomics location-specific barcodes of the array. Using H&E images we estimate each spatial transcriptomics spot captured approximately 20-50 cells. After alignment of fastq file using Cell Ranger (10x Genomics), an average of 3,400 distinct genes were detected in each spot. Each sample was then loaded into R Seurat package and normalized by SCTransform. Clustering was then performed by FindClusters (R Seurat) based on the shared nearest-neighbor modularity optimization algorithm (*30*) using the first 10 dimensions of principal component analysis. Differentially expressed genes with significantly higher expression in each spatial transcriptomics cluster than others were identified (p < 0.01, Wilcoxon rank sum test and average log fold-change > 0) (Supplementary Fig. 1A, B). Publicly available scRNA-seq data from Shih et al. (*34*) were analyzed and used to define differentially expressed genes for the major cell types. For each cell type, genes whose expression was statistically higher in the cells annotated to that cell type in comparison with their expression in the remaining cells were identified (p < 0.00001, Wilcoxon rank sum test, average log_fold-change > 0.25) (Fig. 1B, Supplementary Table 1).

**Fig. 1:**
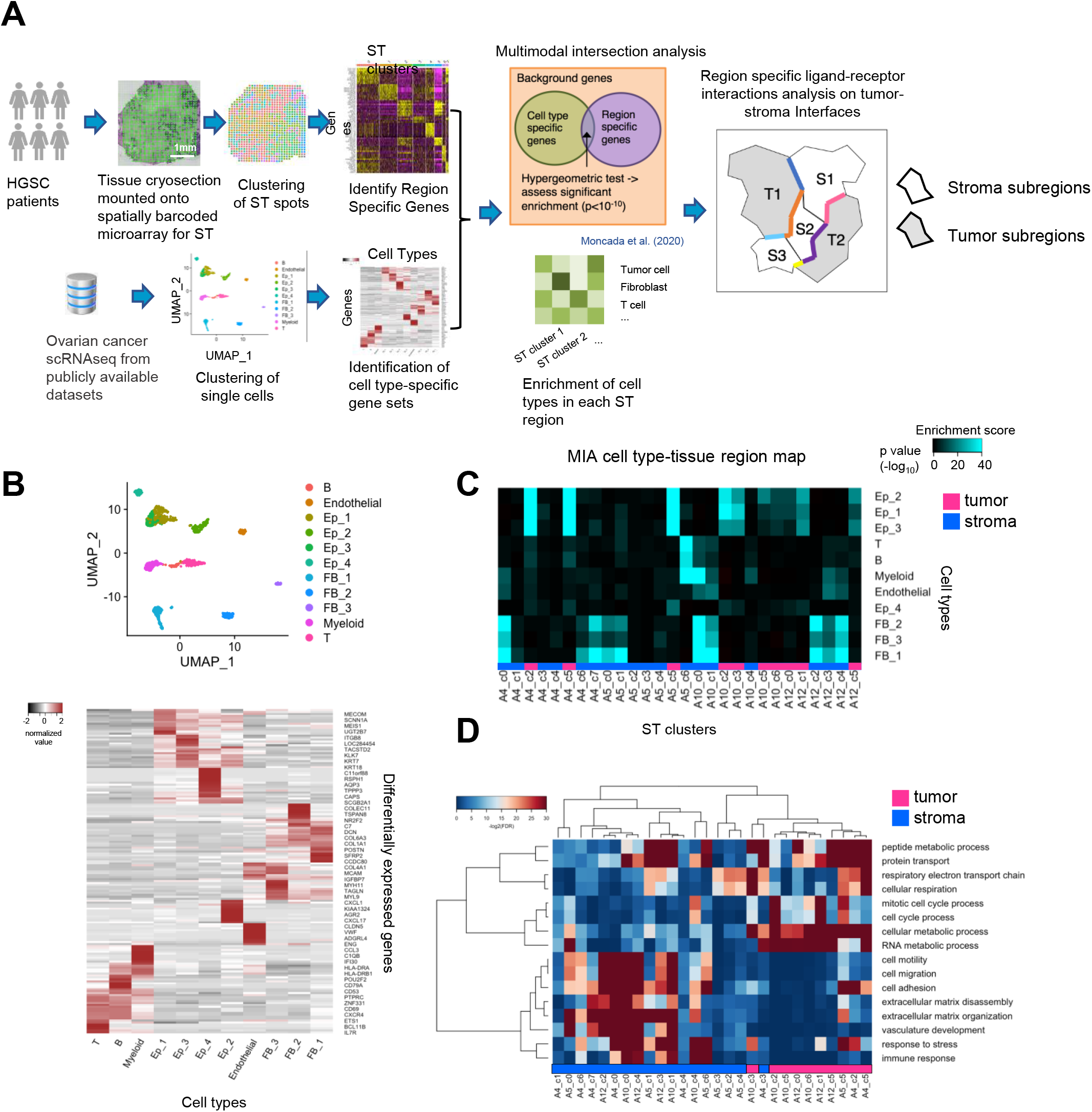
Spatial transcriptomics analysis of four HGSC patient samples. **A.** Schematic of the analysis workflow. The spatial transcriptomics data from our ovarian cancer patient samples were integrated with scRNA-seq data from publicly available datasets. Each spatial transcriptomics tissue cryosection was mounted onto a spatially barcoded microarray. The spots on the microarray were sequenced and clustered based on the gene expression in multiple regions. The scRNA-seq data were also clustered, which generated multiple cell types and their specific gene sets. MIA was performed to identify the overlap between region-specific genes and cell type–specific genes, and a hypergeometric test was performed to determine whether the overlap was significant. The – log_10_(p-value) of the test was used to estimate the enrichment of a cell type in that region. The region-specific ligand-receptor interactions between adjacent stroma and tumor clusters were analyzed. **B.** Top, clustering of scRNA-seq data (n=1,156 cells; data from Shih et al.^31^). Bottom, heatmap of the top 15 differentially expressed genes in each cluster. **C.** The MIA map of all scRNA-seq–identified cell types and spatial transcriptomics-defined regions. Each element in the matrix was computed for all pairs of cell types and tissue regions. The hypergeometric test identified tumor clusters as those enriched with any of the Ep_1, Ep_2, Ep_3, or Ep_4 cell types with a p-value of less than 1 × 10^-10^. A cluster was identified as a stroma cluster if it was not a tumor cluster. The bar at the bottom indicates the region type. **D.** Overrepresentation analysis of the DAVID Gene Ontology gene sets. The bar at the bottom indicates tumor and stroma clusters, which are highlighted in magenta and blue, respectively.

With the differentially expressed genes extracted across the scRNA-seq and spatial transcriptomics modalities, the overlap between each pair of cell type-specific and region-specific gene sets was computed by multimodal intersection analysis (MIA) (*31*). A hypergeometric test was performed to assess significant enrichment using a threshold of p < 10^-10^ (Fig. 1C); for example, tumor clusters were assigned if the enrichment p-value of any of the epithelial cell types Ep_1 (SPON1^+^), Ep_2 (SST^+^), Ep_3 (ATHL1^+^), or Ep_4 (TPPP3^+^) (Supplementary Table 1) in that cluster was lower than 10^-10^. Stroma clusters were assigned if they were not tumor clusters. From the MIA map, we observed the infiltration of immune cells in some tumor clusters (e.g., B cells were enriched in cluster A4_c5 and A4_c2).

Overrepresentation analysis of selected gene ontology (GO) biological process terms showed the difference between tumor and stroma clusters (Fig. 1D and Supplementary Fig. 2). Tumor clusters were highly enriched with metabolic processes (e.g., cellular/RNA metabolism), cellular respiration, and cell cycle processes. Stroma clusters were highly enriched with biological processes of cell motility/migration, extracellular matrix organization/disassembly, vasculature development, and immune responses (Fig. 1D and Supplementary Fig. 2).

### Identification and mapping of tumor and stroma cell clusters

To verify the accuracy of our tumor/stroma classification by enrichment analysis of spatial transcriptomics RNA-seq data, we mapped the normalized expression of major tumor and stroma markers for each sample and overlaid them on the H&E images. We confirmed that the normalized expression of the tumor markers WFDC2, MUC16, and CLDN4 was localized to the tumor regions and that the normalized expression of COL1A1, VIM, and FN1 was restrained to the stroma regions (Fig. 2A and Supplementary Fig. 3).

**Fig. 2.**
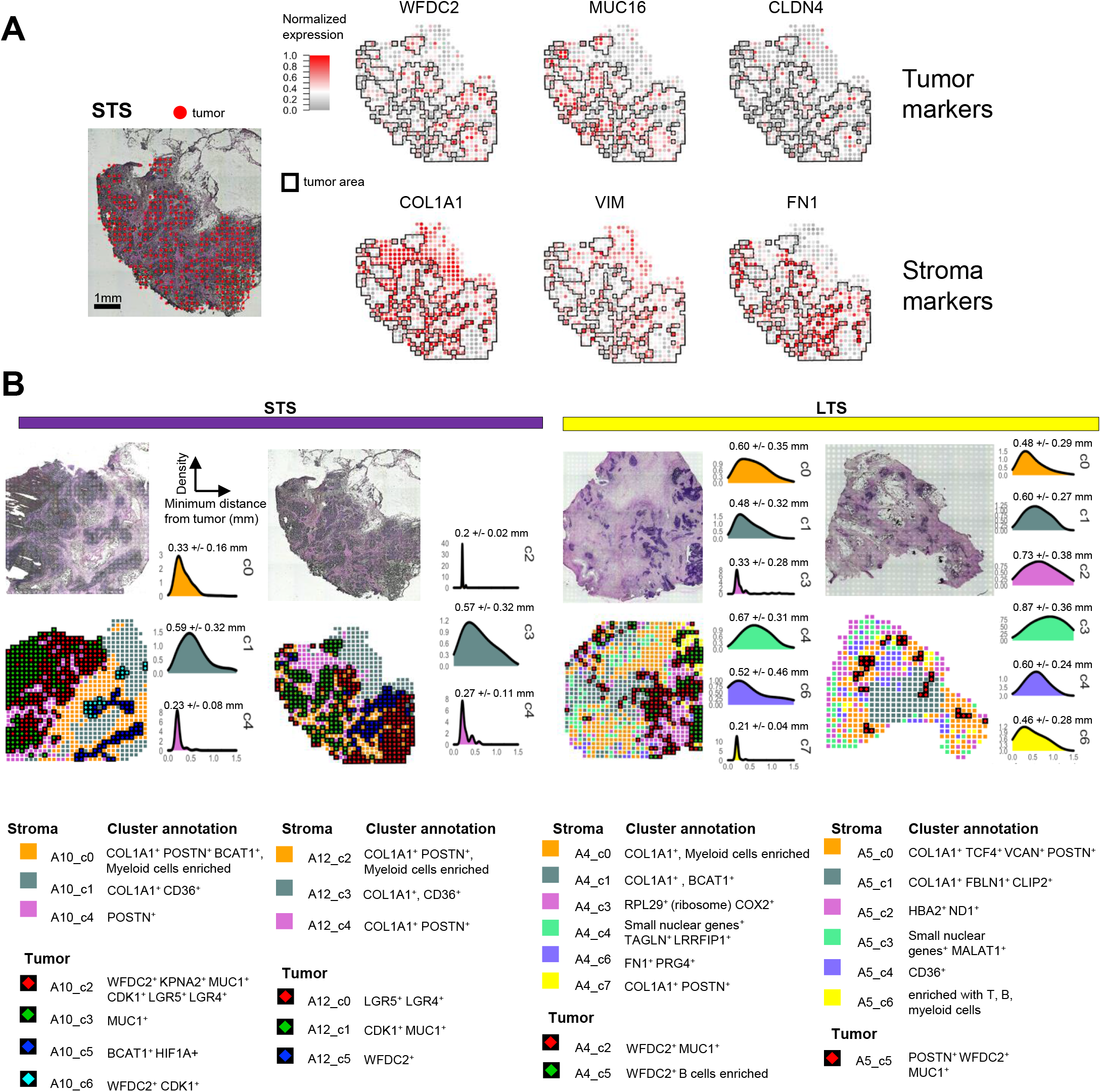
Spatial transcriptomics of HGSC and mapping of clusters. **A.** Spatial expression of marker genes on a representative spatial transcriptomics sample. Left, H&E staining; the tumor spots identified by MIA (red) are overlaid on the tumor regions. Right, normalized spatial expression of the tumor and stroma markers; the black outline indicates the tumor areas overlapping the red spots in the left panel. **B.** Top, H&E staining. Bottom, Clustering of spatial transcriptomics spots. Clusters are annotated by marker genes, the top differentially expressed genes, and MIA enrichment analysis. Samples are grouped as STS samples (left) and LTS samples (right). For each cluster, density plots show the minimum distance from each stroma spot to the tumor. The mean +/- standard deviation for the density distribution is shown at the top of each plot.

After the clustering of spatial transcriptomics spots, we discovered a high level of heterogeneity among the four tumor samples and annotated the tumor and stroma clusters (Fig. 2B). Tumor clusters A10_c2 and A12_c1 (identified in patients A10 and A12) were characterized by the increased expression of CDK1 (avg_logFC = 0.37, p_adj = 1.5*10^-12^ in A10_c2, avg_logFC = 0.31, p_adj = 2.4*10^-5^ in A12_c1, Supplementary Table 2), one of the most important regulators of cell cycle progression in mammalian cells. Moreover, tumor clusters A10_c2 and A12_c0 (in patients A10 and A12, respectively) expressed high levels of LGR5 (avg_logFC = 0.36, p_adj = 2.3*10^-12^ in A10_c2, avg_logFC = 0.49, p_adj = 1.3*10^-17^ in A12_c0, Supplementary Table 2), which promotes cancer cell mobility, tumor formation, and epithelial-mesenchymal transition through the activation of Wnt/β-catenin signaling. In contrast, neither CDK1 nor LGR5 were detected in any of tumor clusters in patients A4 and A5. Moreover, most of the tumor clusters in all four samples expressed WFDC2, a gene commonly overexpressed in ovarian carcinomas compared with normal ovarian tissues (*35*).

### Heterogeneity of stroma subclusters

Analysis of the stroma clusters from patients A10 and A12 showed that the stroma clusters A10_c0 and A12_c4, which were located near the tumor edge, expressed high levels of COL1A1 and periostin (POSTN), whereas stroma clusters A10_c1 and A12_c3, which were located distant from the tumor edge, expressed high levels of COL1A1 and CD36 (Fig. 2B, left panel). Stroma clusters A10_c4 and A12_c2, which were closely surrounded by tumor clusters, expressed high levels of periostin. In contrast, high levels of periostin were identified in only a single stroma cluster, A4_c7, surrounding the tumor cluster A4_c5 in patient A4 and in a single stroma cluster, A5_c0, surrounding the tumor cluster A5_c5 in patient A5. In addition, only stroma cluster A5_c4 in patient A5 expressed CD36 (Fig. 2B, right panel). These data suggest both intra- and inter-stromal heterogeneity among HGSCs. Moreover, tumors derived from patients A10 and A12 were distinct from those derived from patients A4 and A5. Indeed, survival analysis demonstrated that patients A4 and A5 were long-term survivors (LTSs) with overall survival durations of more than 120 months, whereas patients A10 and A12 were short-term survivors (STSs) with overall survival durations of less than 24 months. Furthermore, tumor samples from LTSs had lower levels of stromal periostin and CD36 expression. These finding suggest that low levels of periostin and CD63 expression in the stroma compartment of the tumor tissue may serve as prognostic markers of long-term survival.

To further elucidate stromal heterogeneity and the relationship between traditional CAF markers and the stroma clusters, we assessed the expression of traditional CAF markers, including αSMA (ACTA2), S100A4, VIM, FAP, PDGFRα (PDGFRA), and PDGFRβ (PDGFRB), in the stroma clusters we identified. Because we demonstrated that stromal CD36 and periostin were associated with STS (samples A10, A12) (Fig. 2B), which indicated a more aggressive phenotype of the stroma clusters, we included these markers in our CAF gene panel. Only 2 of the 6 stroma clusters in both the LTS samples (A4 and A5) expressed the traditional CAF markers, whereas all the stroma clusters in the STS samples (A10 and A12) expressed the CAF markers (Fig. 3A). These findings suggest these stroma clusters are enriched with CAFs in the tumor tissue and only a small percentage of stroma clusters in LTS tumors are enriched with CAFs. The stroma clusters that did not express CAF markers may be enriched with quiescent fibroblasts, mesenchymal cells, or other stromal cell types. In the two STS samples, the CAF clusters expressed a different set of CAF markers, and most expressed a majority of the eight CAF markers. In contrast, in the two LTS samples, all the stroma clusters that had CAF marker expression expressed a minority of the eight CAF markers. (Fig. 3A-C and Supplementary Fig. 4). These findings suggest intra- and inter-tumor CAF heterogeneity in HGSC and that tumors with enriched CAF clusters expressing multiple CAF markers may be associated with short-term survival.

**Fig. 3.**
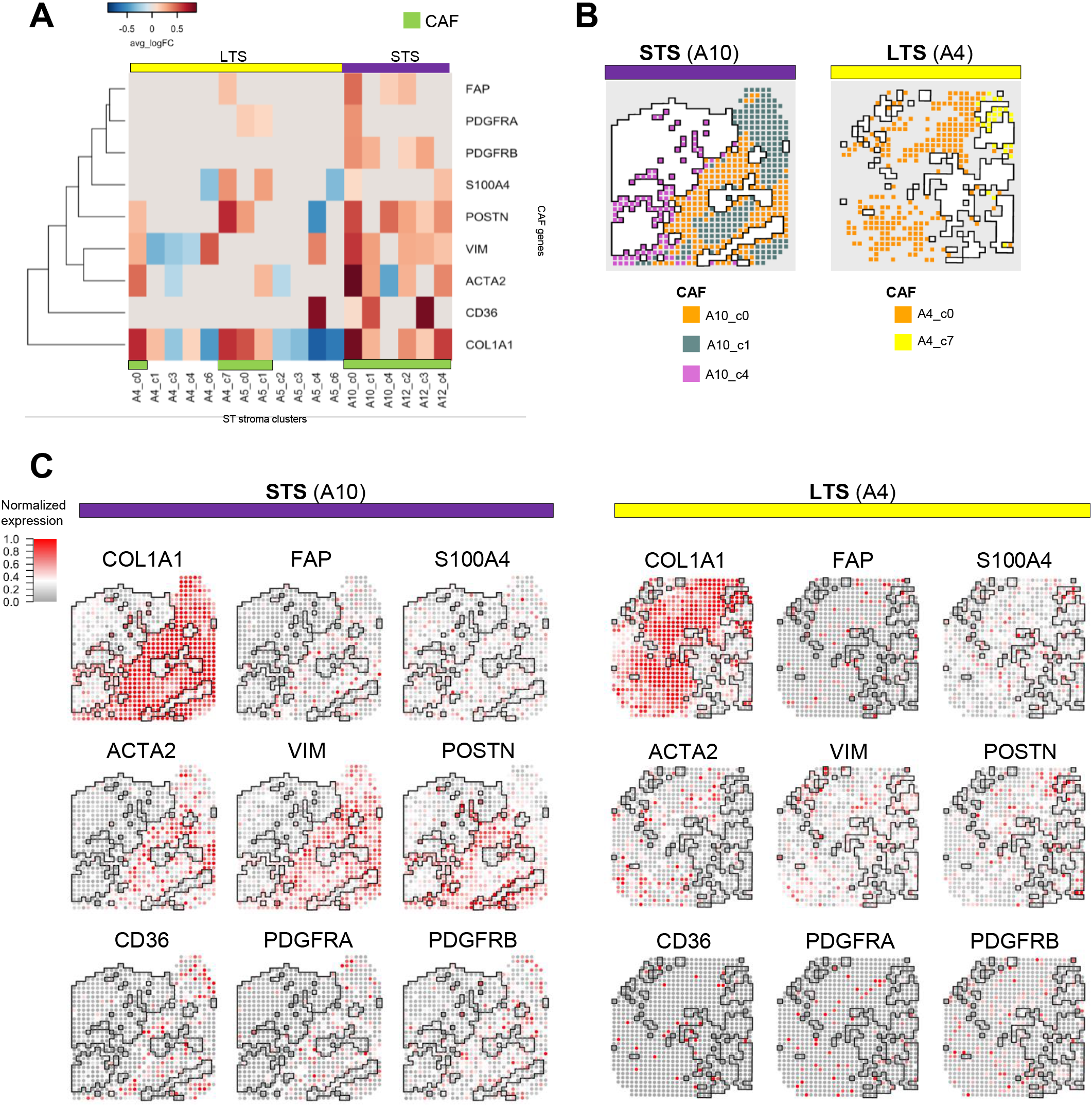
Heterogeneity of CAFs. **A.** Heatmap of the average log fold-change of CAF gene expression in the stroma clusters. The average log fold-change was computed as the log fold-change of the average expression of a gene between a cluster and all the other clusters of the sample. The average log fold-change was set to 0 if the p-value was larger than 0.01 (Wilcoxon rank sum test). **B.** Mapping of CAF clusters in an STS sample (left) and an LTS sample (right). **C.** Spatial expression of selected CAF genes in an STS sample (left) and an LTS sample (right).

Next, we examined the spatial distribution of various CAF clusters and the key molecules used for their annotation relative to the tumor clusters. The average expression of CAF markers as a distance to tumor showed that the overall CAF localization relative to tumor varied across samples and different among CAF markers (Supplementary Fig. 4C). CAF clusters in the neighborhood of the tumor clusters (e.g., A10_c0, A10_c4, and A4_c7) expressed high levels of periostin (Fig. 3B, C and Supplementary Fig. 4). Furthermore, CAF clusters located distant from the tumor clusters (e.g., A10_c1 and A5_c4) expressed high levels of CD36. These findings suggest that spatially resolved CAF subpopulations in the ovarian TME may play different roles in conferring the malignant phenotypes of the tumor cells.

### Prognostic significance of CD36 and periostin

Because the spatial transcriptomics data demonstrated that the overall periostin and CD36 expression levels in the CAF clusters of the 2 STS samples were markedly higher than those in the 2 LTS samples, we performed multiplexed immunofluorescence (mIF) analysis to quantify the spatial expression of periostin and CD36 in an independent set of 27 advanced HGSC cases (11 LTS and 16 STS) to verify our findings. Measuring the mean intensity of the selected markers in only the stroma areas demonstrated that STS samples had significantly higher periostin expression levels near the stroma-tumor interface than LTS samples did (p = 0.0063) (Fig. 4A, C). In addition, CD36 expression levels in the stroma areas distant from the tumor clusters were markedly higher in STS samples than they were in LTS samples, but this difference was not statistically significant (p = 0.0575) (Fig. 4B, D).

**Fig. 4.**
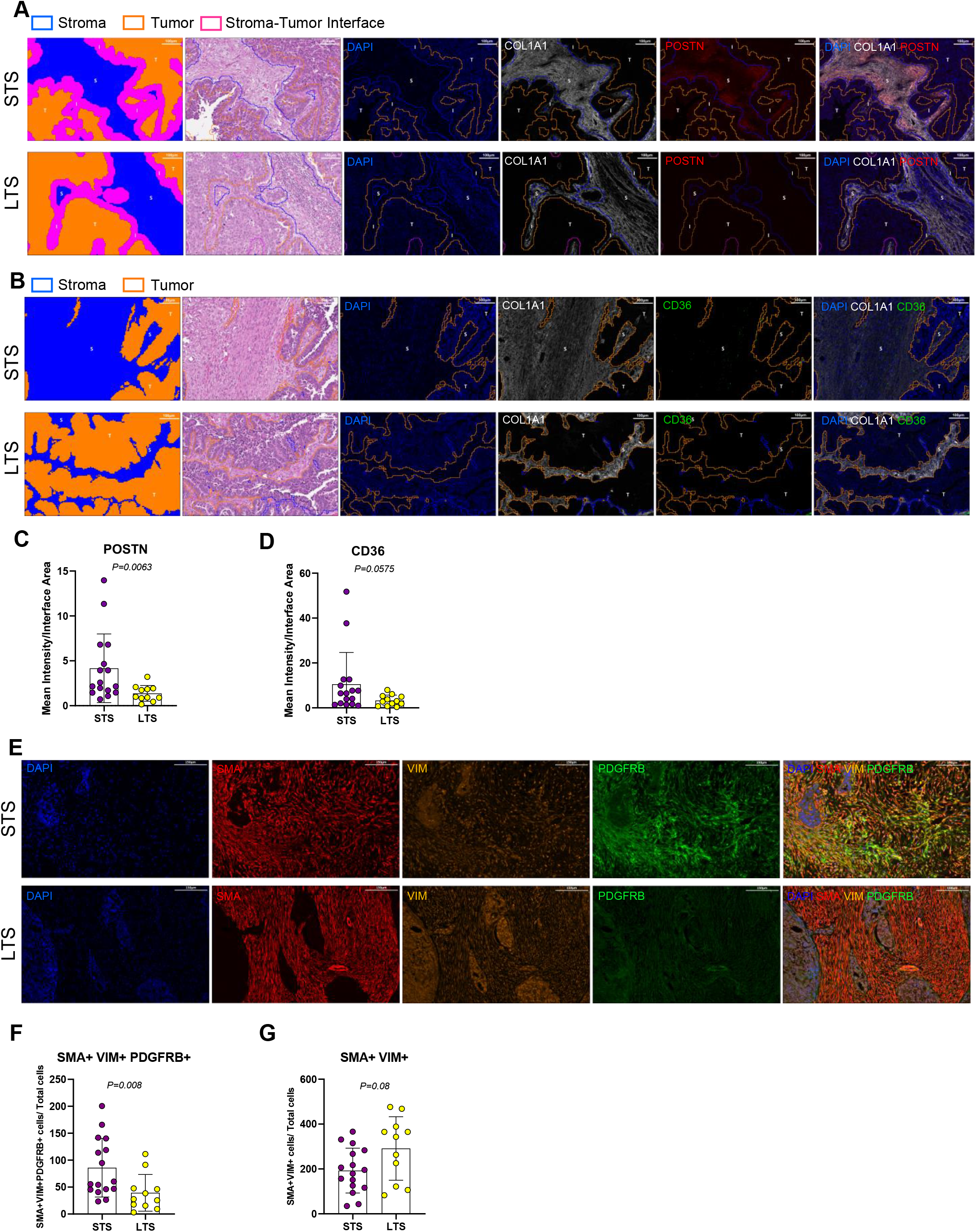
Validation of CAF subclusters. **A-D.** mIF staining of markers of stroma subclusters in paraffin-embedded HGSC tissue: periostin (POSTN; A and C) and CD36 (B and D). Scale bar for panel A is 100μm. **E-G.** mIF staining of markers of CAF subclusters. Scale bar for panel E is 150μm.

To validate the presence of a unique set of CAFs in STS samples compared to LTS samples and to validate our spatial transcriptomics findings showing that STS samples have markedly increased CAF clusters expressing all three markers (VIM, αSMA, and PDGFRB), we performed mIF analysis of these markers. We calculated the density of typical CAFs, expressing VIM, αSMA, and PDGFRB, by normalizing their number by the total number of stromal cells. We found a significantly higher density of αSMA^+^VIM^+^PDGFRβ^+^ cells in the stroma of STS samples than in the stroma of LTS samples (p = 0.008) (Fig. 4E, F). The density of αSMA^+^VIM^+^ cells, which represent myofibroblasts, did not differ between the STS and LTS samples (p = 0.08) (Fig. 4E, G). These results indicate that the spatially resolved CAF subpopulations expressing POSTN and αSMA, VIM, and PDGFRB in the ovarian TME play a major role in conferring a more aggressive type of HGSC with decreased overall survival.

### Crosstalk signaling network analysis

To characterize region-specific ligand-receptor interactions between CAFs and tumor cells, crosstalk signaling network analysis was performed. For each tumor and CAF cluster in each sample, we selected only the nearest neighboring spots on the interface and performed ligand-receptor analysis (Fig. 5A). In total, 84 and 50 distinct ligand-receptor pairs at the stroma-tumor interfaces of STS and LTS samples were identified, respectively (Supplementary Fig. 5-7). For example, we found APOE (ligand; on CAF) and LRP5 (receptor; on tumor) at the A10_c0-A10_c2 interface and the A12_c4-A12_c5 interface and found THBS2 (ligand; on CAF) and CD47 (receptor; on tumor) at the A10_c0-A10_c6 interface (Fig. 5B,C and Supplementary Fig. 8). The region-specific ligand-receptor interaction networks between adjacent stroma and tumor subregions are shown in Fig. 5D. Sixty distinct ligand-receptor pairs were identified and pooled from the stroma-tumor interfaces between every stroma and tumor cluster in sample A10 (Fig. 5E). By matching this information with published data (searching for correlation with overall survival worst prognosis, and metastatic process), we identified 9 ligand-receptor pairs associated with STS samples (APOE-LRP5, THBS2-CD47, PLAU-PLAUR, WNT10A-FZD5/FZD7, TGFB1-ENG/ACVRL1, IGF1-IGF1R, SEMA3C-NRP2) and 1 ligand-receptor pair associated with LTS samples (A2M-LRP1) (Fig. 5B, C and Supplementary Fig. 8).

**Fig. 5.**
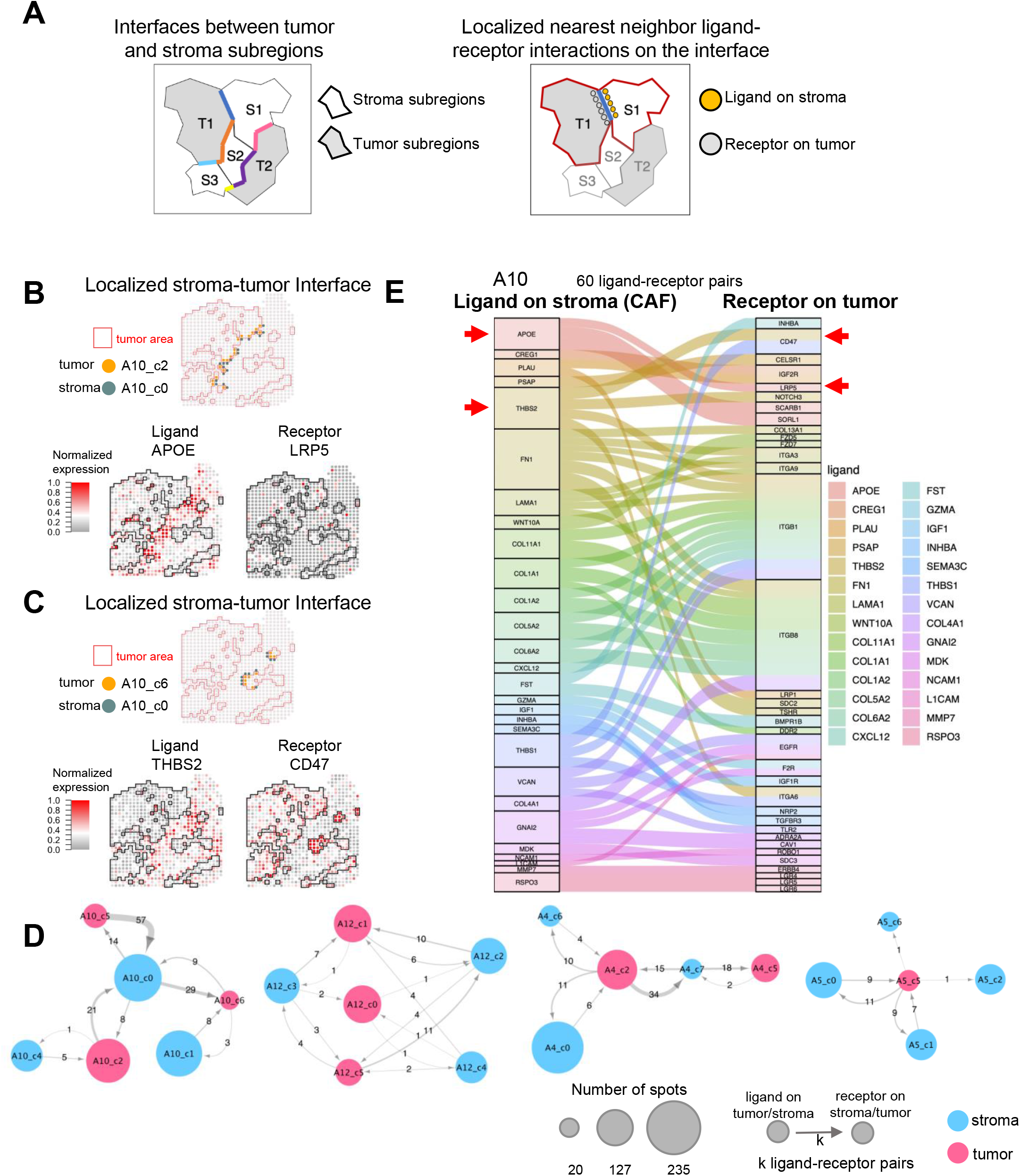
Region-specific ligand-receptor interactions between stroma and tumor. **A.** Illustration of the method used to identify the region-specific ligand-receptor interactions. Left, Different stroma-tumor interfaces in a sample. Right, the method identifies the ligand-receptor pairs at each interface. **B, C.** Representative nearest-neighbor interactions between tumor and stroma subregions. Top, Localization of nearest-neighbor spots of the tumor subregion (orange) and stroma subregion (gray). Red outlining indicates the tumor area of the sample. Bottom, the expressions of the ligand (left)-receptor (right) pairs are localized near the interface shown in the top panel. **D.** Nearest-neighbor interaction networks between tumor and stroma subregions in all four samples. Pink and blue indicate tumor and stroma subregions, respectively. The node size is proportional to the number of spots in a subregion, and the width of the edges between nodes is proportional to the number of ligand-receptor pairs. **E.** Alluvial plot of all ligand-receptor pairs between tumor and stroma in a representative sample. Sixty ligand-receptor pairs were identified and pooled from all the interfaces between every stroma and tumor subregion.

Because the ligand-receptor pair APOE-LRP5 was found in the subcluster interface of both STS samples, we used mIF analysis to validate the presence of this interaction in tumor samples obtained from the same cohort of LTS patients (n=11) and STS patients (n=16) used for CD36 and periostin staining as described above. Tissue samples were stained for COL1A1, LRP5, and APOE. The number of LRP5^+^ cells and the mean intensity of APOE expression was calculated at the stroma-tumor interface. Spearman correlation analysis revealed a significant positive correlation between LRP5^+^ cell density and APOE expression intensity at the stroma-tumor interface of STS samples (r = 0.6, p = 0.0261) (Fig. 6A, B) but not LTS samples (r = −0.2484, p = 0.3911) (Fig. 6A, C). Moreover, no correlation between LRP5^+^ cells in tumor areas and APOE expression intensity in stroma areas (excluding the stroma-tumor interface area) in either STS or LTS samples was found (r = 0.4505, p = 0.1081 and r = −0.13141, p = 0.6485, respectively) (Supplementary Fig. 9A). Furthermore, there was no significant correlation between APOE expression intensity and LRP5^+^ cells in either the tumor or stroma areas of STS samples (r = 0.02857, p = 0.9276 and r = 0.4505 and p = 0.1081, respectively) or LTS samples (r = −0.2967, p = 0.3025 and r = 0.03736 and p = 0.9035, respectively) (Supplementary Fig. 9B, C). These data suggest that significant APOE-LRP5 crosstalk occurs at the stroma-tumor interface only and that such crosstalk plays a crucial role in modulating the malignant phenotype of HGSC, which could serve as a predictive biomarker of patient survival.

**Fig. 6.**
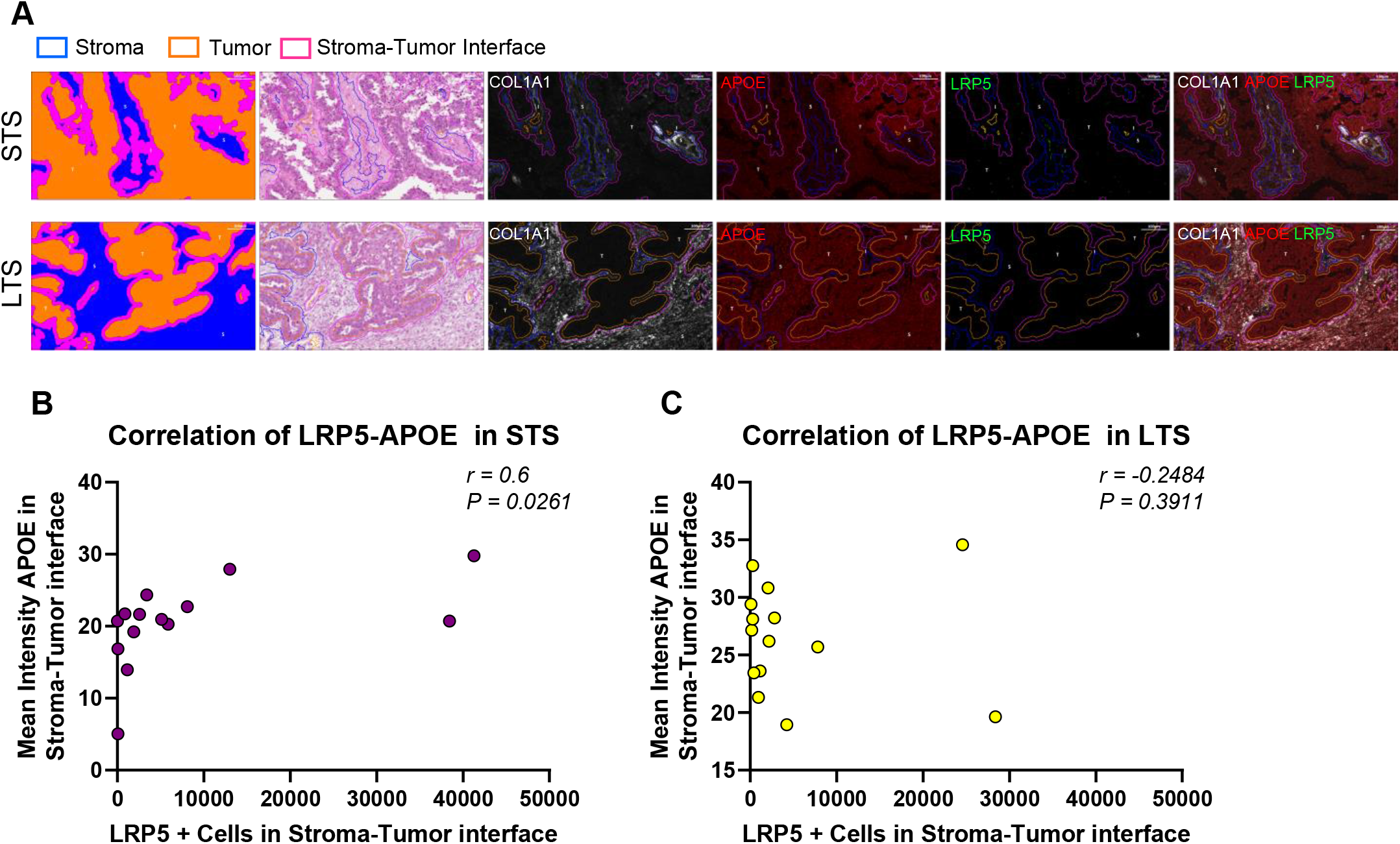
Validation of ligand-receptor cross-talk at the stroma-tumor interface. **A.** mIF staining of the ligand-receptor pair APOE-LRP5 in paraffin-embedded HGSC tissues. **B, C.** Spearman’s correlation analysis between LRP5^+^ cells and APOE expression intensity in the stroma-tumor interface area in STS samples (B) and LTS samples (C).

## DISCUSSION

In this study, spatial transcriptomics technology was used to characterize CAF heterogeneity in advanced-stage HGSC. We found the absence of specific CAF subtypes in the tumor tissue of patients with advanced HGSC to be associated with survival durations of more than 10 years. Furthermore, specific ligand-receptor interactions between various tumor and CAF clusters in STS samples compared with those in LTS samples were identified.

Although most ovarian cancer patients have a median survival duration of less than 5 years, approximately 15% of patients survive more than 7 years (*6*). These patients are generally defined as long-term survivors (LTSs) (*36*). Clinical features, including histological grade and type, age at diagnosis, and optimal cytoreduction at primary surgery, are associated with long-term survival (*5*). In previous studies, multiple transcriptome analyses of LTS samples were performed to identify conserved genomic signatures associated with long-term survival; however, each study identified different gene sets, most likely because of the use of heterogenous patient cohorts and the use of bulk tissue samples with various amount of stromal tissue (*21*). The gene sets identified may not represent the biology of the tumor that contribute to long-term survival. In the present study, to eliminate the confounding factors associated with patient survival, we included only optimally debulked, advanced-stage, treatment-naïve HGSC samples obtained from STSs (overall survival < 24 months) and LTSs (overall survival > 120 months). The use of a spatially resolved transcriptomic platform can also eliminate the issue of tissue heterogeneity among different specimens.

Our spatial transcriptomics analysis demonstrated the presence of different tumor clusters represented by different transcriptome signatures within a tumor tissue, suggesting high levels of heterogeneity not only among different HGSC samples but also within single HGSC samples. For example, the annotated A10_c2 and A12_c1 tumor clusters and A10_c2 and A12_c0 tumor clusters identified in STS but not in LTS samples expressed high levels of CDK1 and LGR5, respectively, which suggests that a cluster of tumor cells with CDK1 or LGR5 expression in the tumor tissue of an STS sample expressing one of these genes may play a role in conferring the malignant phenotype of HGSC in STSs. Indeed, several studies have shown that CDK1 dysregulation leads to robust tumor growth, chromosomal instability, and a high rate of tumor cell proliferation (*37*). Furthermore, other studies have demonstrated that LGR5 expression levels are positively correlated with cancer stem cell traits, shorter survival times, and chemoresistance (*38, 39*).

Increasing evidence suggests that the stromal component of the TME plays an important role in ovarian cancer development. In HGSC patients, the high number of stromal cell types in the ovarian TME is significantly associated with poor survival (*40*). In contrast to lymphocytic infiltration, which is correlated with favorable prognosis, the presence of specific types of CAFs can predict the overall survival and progression-free survival of HGSC patients (*40, 41*). Recently, Stur et al. used spatial transcriptomics to identify spatially resolved biomarkers that can be used to differentiate between HGSC patients whose disease has a poor or excellent response to chemotherapy (*42*). They demonstrated the importance of the stromal component and the presence of different clusters as drivers of poor response (*42*). However, the clusters they identified were not annotated, and the expression of marker genes in the clusters were not validated at the protein level. Nevertheless, the authors demonstrated the feasibility of using spatial transcriptomics to identify predictive markers of treatment response. In the present study, our spatial transcriptomics analysis of treatment-naïve STS and LTS samples demonstrated high levels of both intra- and inter-stromal heterogeneity among HGSCs. Annotated stroma clusters, which were identified predominantly in STS samples, had high levels of both periostin and CD36 expression as well as high levels of COL1A1 expression. We were able to validate these findings in a larger cohort of patients using mIF analysis, which revealed a significantly higher level of periostin protein expression in a specific stroma compartment of STS samples than in that of LTS samples. Markedly higher levels of CD36 protein expression were also observed in STS; however, this difference was not significant. Periostin, a component of the extracellular matrix, is expressed by fibroblasts in normal tissue and by those in the stroma of various primary tumors. Periostin is also required for cancer stem cell maintenance, and blocking its function prevents metastasis (*43*). Periostin is also associated with poor prognosis and platinum resistance in epithelial ovarian cancer (*44*). CD36 fuels tumor metastasis and therapy resistance by enhancing lipid uptake and fatty acid oxidation. It also attenuates angiogenesis by binding to thrombospondin 1 and thereby inducing apoptosis or blocking the VEGFR-2 pathway in tumor and endothelial cells. Moreover, CD36-driven lipid metabolic reprogramming and functions in tumor-associated immune cells lead to tumor immune tolerance and cancer development (*45*). Therefore, the overexpression of periostin and CD36 in the stromal compartment could contribute to the malignant phenotype of HGSC in STS.

To further investigate whether the stromal heterogeneity we identified represents different CAF populations, we assessed the expression of canonical CAF markers in the stroma clusters we identified. Stroma clusters in STS samples expressed all traditional CAF markers, including αSMA (ACTA2), S100A4, VIM, FAP, PDGFRα, and PDGFRβ, which suggests that these clusters represent CAF clusters. However, most stroma clusters in LTS samples expressed only αSMA and VIM but no other CAF markers, which suggests that these clusters represent myofibroblasts rather than canonical CAFs. Validation studies using mIF confirmed that the density of CAFs expressing αSMA, VIM, and PDGFRB in LTS samples was significantly lower than that in STS samples, indicating that the absence of CAF clusters that express all the canonical CAF markers can be used to predict long-term survival.

Because the crosstalk between CAFs and neighboring tumor cells plays an important role in modulating the malignant phenotype of cancer, we characterized region-specific ligand-receptor interactions by analyzing crosstalk signaling in neighboring spots at the stroma-tumor interface. Crosstalk analysis methods have been designed for use with both cell-specific RNA-seq data (e.g., CCCexplorer (*32*)) and single-cell RNA-seq data (*46, 47*). For spatial transcriptomics data, however, there is a lack of systems approaches for analyzing the cell-cell crosstalk. In this paper, we propose a method for studying region-specific ligand-receptor interactions at the stroma-tumor interface. The method can also be used to analyze crosstalk between other cell types (e.g., between tumor cells and immune cells, between cells in the immune stroma, between different tumor cells). We selected receptors and ligands in tumor spots and stroma spots, respectively, at the interface and identified 9 and 1 ligand-receptor pairs at the interfaces in STS and LTS samples, respectively. Because the APOE-LRP5 ligand-receptor pair was identified by spatial transcriptomics in both STS samples, we used mIF to assess the correlation between LRP5^+^ cells and the mean intensity of APOE expression in a larger cohort of HGSC samples. We found a significant positive correlation between LRP5^+^ cells and APOE intensity in only STS samples in which this ligand-receptor pair is exclusively expressed at the stroma-tumor interface. This suggests a strong crosstalk signaling network between tumor-derived LRP5 and CAF-derived APOE, which may contribute to poor survival. Increased levels of LRP5 have been linked to metastasis in various tumor types (*48–50*). Besides being involved in the Wnt/βcatenin canonical pathway, LRP5 has been reported to be involved in the uptake of glucose in mammary epithelial cells through APOE binding, which is essential for regulating the growth rate of these cells (*50*). Moreover, APOE has been reported to be required for cell proliferation and survival in ovarian cancer and has been associated with aggressive biology and poor prognosis in colorectal cancers (*51, 52*).

One of the present study’s limitations was that only 4 samples (2 STS samples and 2 LTS samples) were used in the initial spatial transcriptomics analysis. However, mIF analysis in a larger cohort of independent samples validated our findings, suggesting that our analytical methods were robust and stringent. The study had a small sample size owing to the rarity of LTSs with advanced HGSC, which comprise only 15% of HGSC patients (*6*), and its use of treatment-naïve samples. The use of spatial transcriptomics platforms with a more homogenous patient cohort will certainly increase the likelihood of identifying reliable prognostic or predictive biomarkers of advanced HGSC.

In conclusion, our spatial transcriptomics analysis revealed high levels of inter- and intra-tumor CAF heterogeneity as well as novel spatially resolved CAF-tumor crosstalk signaling networks in the ovarian TME that are associated with long-term survival in patients with advanced HGSC. Further elucidating the association between these spatially resolved biomarkers and long-term survival could contribute to our understanding of the biology of ovarian cancer and thereby improve the survival of patients with advanced HGSC.

## ACKNOWLEDGMENTS

We are grateful for the generous donation of tissue samples by patients undergoing surgery. We thank Siu Fee Rita and Mansi Diwanji for excellent technical support with mIF slide staining. We are also grateful to Joseph Munch from the Scientific Publications, Research Medical Library at MD Anderson Cancer Center for help editing the manuscript.

## AUTHOR CONTRIBUTIONS

S.F.-B., Y.Z., and S.C.M., designed the study and planned the experiments. S.F.-B. conducted the majority of the experiments and analyzed the mIF data and Y.Z. analyzed the ST and scRNA-seq data. J.S., J.K.B., J.G., K.K.W. and S.T.C.W., provided intellectual contributions to experimental design and data analysis. S.F.-B. and Y.Z., wrote the initial draft of the manuscript. S.F-B., Y.Z., S.T.C.W., and S.C.M., prepared and revised the manuscript. All authors have read and agreed to the published version of the manuscript. The authors declare that they have no competing interests.

