## Supplementary Methods and Figures for "Long-term ovarian cancer survivors: spatial transcriptomics depict ligand-receptor crosstalk heterogeneity at the tumor-stroma interface"

##### scRNA-seq analysis

HGSC scRNA-seq datasets were downloaded from the GEO database (see data availability, below). The unique molecular identifier (UMI) count matrices of the cells were loaded into Seurat 3.2 for downstream analyses. Only single-cell transcriptomes with more than 500 UMIs and less than 10,000 UMIs and with less than 25% mitochondrial transcripts were kept. Genes expressed in less than two cells were removed. UMI counts were normalized by the total counts, multiplied by a default scale factor of 10,000, and then log-transformed. This was followed by principal component analysis of the top 3,000 most variable genes in the dataset; clustering using a shared nearest-neighbor (SNN) modularity-based clustering algorithm (the Seurat default clustering method) (*1*), which used the top 10 components of the principal component analysis (PCA) to find neighbors; and dimensionality reduction for visualization using uniform manifold approximation (UMAP). We defined genes differentially expressed between one cluster and all other clusters as those with a p-value less than 0.00001 (Wilcoxon rank sum test) and an average log fold-change greater than 0.25. The average log fold-change was computed as the $\log_e$ fold-change of the average expression level of a gene between a cluster and all other clusters of the sample. Only the genes that were detected in at least 25% of the cells in either of the two populations were kept.

The scRNA-seq dataset had 1,156 cells from HGSC patients that passed the UMI thresholding described above. The clustering process generated 11 clusters. The cell type–specific markers identified among the differentially expressed genes of each cluster were used to determine the putative identities of each cell cluster. For example, CD3E, CD3D, and CD2 were T-cell markers; CD79A, BANK1, and MS4A1 (CD20) were B-cell markers; EMCN and PECAM1

(CD31) were endothelial cell markers; LILRB4, ITGAM (CD11b), and CD163 were myeloid cell markers; COL1A1, ACTA2, and DCN were fibroblast markers; and EPCAM, KRT8, KRT18, WFDC2, and MUC1 were epithelial cell markers. Based on these cell-type specific markers, these 11 clusters were annotated as epithelial cells (clusters Ep_1 [215 cells], Ep_2 [126 cells], Ep_3 [108 cells], and Ep_4 [57 cells]), fibroblasts (clusters FB_1 [150 cells], FB_2 [88 cells], and FB_3 [41 cells]), endothelial cells (56 cells), B cells (47 cells), T cells (149 cells), and myeloid cells (119 cells).

**Spatial transcriptomics RNA-seq analysis**

After the alignment, filtering, barcoding, and UMI counting of the fastq files using Cell Ranger (10x Genomics), an average of 3,400 distinct genes were detected in each spot. Spots with less than 200 genes and genes expressed in less than 2 spots in a sample were removed. The output files were loaded into the Seurat package in R and normalized with SCTransform. The spatial transcriptomics spots were clustered by the same method used in the scRNA-seq data analysis, in which the top 10 components of the PCA were used to find neighbors. Genes differentially expressed between one spatial transcriptomics cluster and all other clusters were defined as those with a p-value less than 0.01 (Wilcoxon rank sum test) and an average log fold-change greater than 0.

**Gene Ontology overrepresentation analysis**

We performed a hypergeometric test to estimate the enrichment of the GO terms in each spatial transcriptomics cluster by computing the overlap between any GO term gene set and the differentially expressed genes of each spatial transcriptomics cluster (average log fold-change >

0, $p < 0.01$). The number of background genes was set at 20,000. The top 10 GO terms for each cluster were kept. Only the GO terms with a false discovery rate (FDR) of less than 0.01 in at least one cluster were kept. Data were displayed using heatmaps in which the rows and/or columns were arranged by hierarchical clustering with (1-spearman_corr) as the distance metric.

**Ligand receptor interaction analysis**

To identify the ligand-receptor pairs in which the ligand was on stroma and the receptor was on tumor, we identified the differentially expressed ligands ($p < 0.05$, average $\log_e$ fold-change > 0.25) that occurred in more than 10% of stroma spots near an interface by comparing the ligand gene expression of the stroma spots near an interface to all the other stroma spots (including spots in other clusters) and identified the differentially expressed receptors ($p < 0.05$, average $\log_e$ fold-change > 0) that occurred in more than 10% of tumor spots near an interface by comparing the receptor gene expression of the tumor spots near an interface to all the other tumor spots (including spots in other clusters). We used a similar approach to identify the ligand-receptor pairs in which the ligand was on tumor and the receptor was on stroma. We identified 84 and 50 distinct ligand-receptor pairs present at the stroma-tumor interface in STS and LTS samples, respectively. We used the ggalluvial package in R to create alluvial plots of the ligand-receptor pairs and used Cytoscape 3.7 (Cytoscape, RRID:SCR_003032) to create figures of nearest-neighbor interaction networks. The connection weight is proportional to

$$\sqrt[4]{\text{avg\_logFC(ligand)} * \text{avg\_logFC(receptor)} * \text{fraction\_ligand} * \text{fraction\_receptor}},$$

where fraction_ligand and fraction_receptor are the percentages of interface spots for which the ligand gene count was greater than 0 and the receptor gene count was greater than 0, respectively.

**Multiplexed immunofluorescence**

A total of 27 paraffin-embedded tumor tissue samples obtained from patients with treatment-naïve advanced HGSC were stained with a panel of 3 antibodies: anti-COL1A1 (Cell Signaling Technology, Cat# 66948, RRID:AB_2920541), anti-periostin (R and D Systems Cat# AF3548, RRID:AB_2166755), and anti-CD36 (Sigma-Aldrich Cat# HPA002018, RRID:AB_1078464). Alternatively, tissue samples were stained with antibodies against αSMA (Cell Signaling Technology Cat# 56856, RRID:AB_2799522), VIM (Abcam Cat# ab11256, RRID:AB_2241674), and PDGFRβ (Cell Signaling Technology Cat# 3169, RRID:AB_2162497). For ligand-receptor crosstalk validation, tissue samples were stained with antibodies against COL1A1 (Cell Signaling Technology, Cat# 66948, RRID:AB_2920541), LRP5 (Thermo Fisher Scientific Cat# PA5-18387, RRID:AB_10979335), and APOE (Millipore Cat# AB947, RRID:AB_2258475). After overnight incubation with the primary antibodies at 4°C, tissue slides were then incubated with a mixture of fluorescent-labeled secondary antibodies: donkey anti-goat Alexa Fluor 594 (Thermo Fisher Scientific Cat# A32758, RRID:AB_2762828), donkey anti-rabbit Alexa Fluor 488 (Thermo Fisher Scientific Cat# A32790, RRID:AB_2762833), and donkey anti-mouse Alexa Fluor 647 Thermo Fisher Scientific Cat# A32787, RRID:AB_2762830). Slides were then counterstained with DAPI (Thermo Fisher Scientific) for nuclei detection. Whole-tissue slides were then scanned utilizing the Vectra Polaris multispectral imaging system (PerkinElmer) at MD Anderson's Flow Cytometry and Cellular Imaging Facility. Tissue slides were then counterstained with H&E and imaged again. Details of the image analysis are provided in the Supplementary Data.

Image analysis was performed using the Visiopharm image analysis software. Fluorescent and colorimetric images of each sample were first aligned to obtain a 3-dimensional image. The H&E layer was used for tissue segmentation to separate tumor and stroma areas, and the tumor and stroma areas were each eroded by 25 μm to obtain an interface width of approximately 50 μm. The mean intensity of CD36 and periostin expression in the stroma areas was then calculated for each sample and normalized by the total stroma area/sample. For the tissue sections stained for αSMA, VIM, and PDGFRβ, tissue segmentation based on αSMA expression was performed to separate tumor and stroma areas. Cell boundaries were determined by a pretrained machine learning algorithm that that used DAPI channel to automatically identify nuclei and cells. Identified cells were then phenotyped using Visiopharm's unbiased autoclustering module using only the top 20% of pixel values per cell. The number of positive cells was then calculated and normalized by the total number of cells in the stroma area/sample. To validate ligand-receptor crosstalk, we performed tissue segmentation and identified the stroma-tumor interface as described above. Densities of cells with LRP5 expression (detected by the 488 channel) were determined by the machine learning algorithm as described above, and the mean intensity of APOE expression (detected by the 594 channel) in the tumor and stroma areas and at the stroma-tumor interface was calculated. Spearman correlation was then performed to determine the extent to which $LRP5^{+}$ cell density and mean APOE expression intensity were correlated.

### SUPPLEMENTARY FIGURES

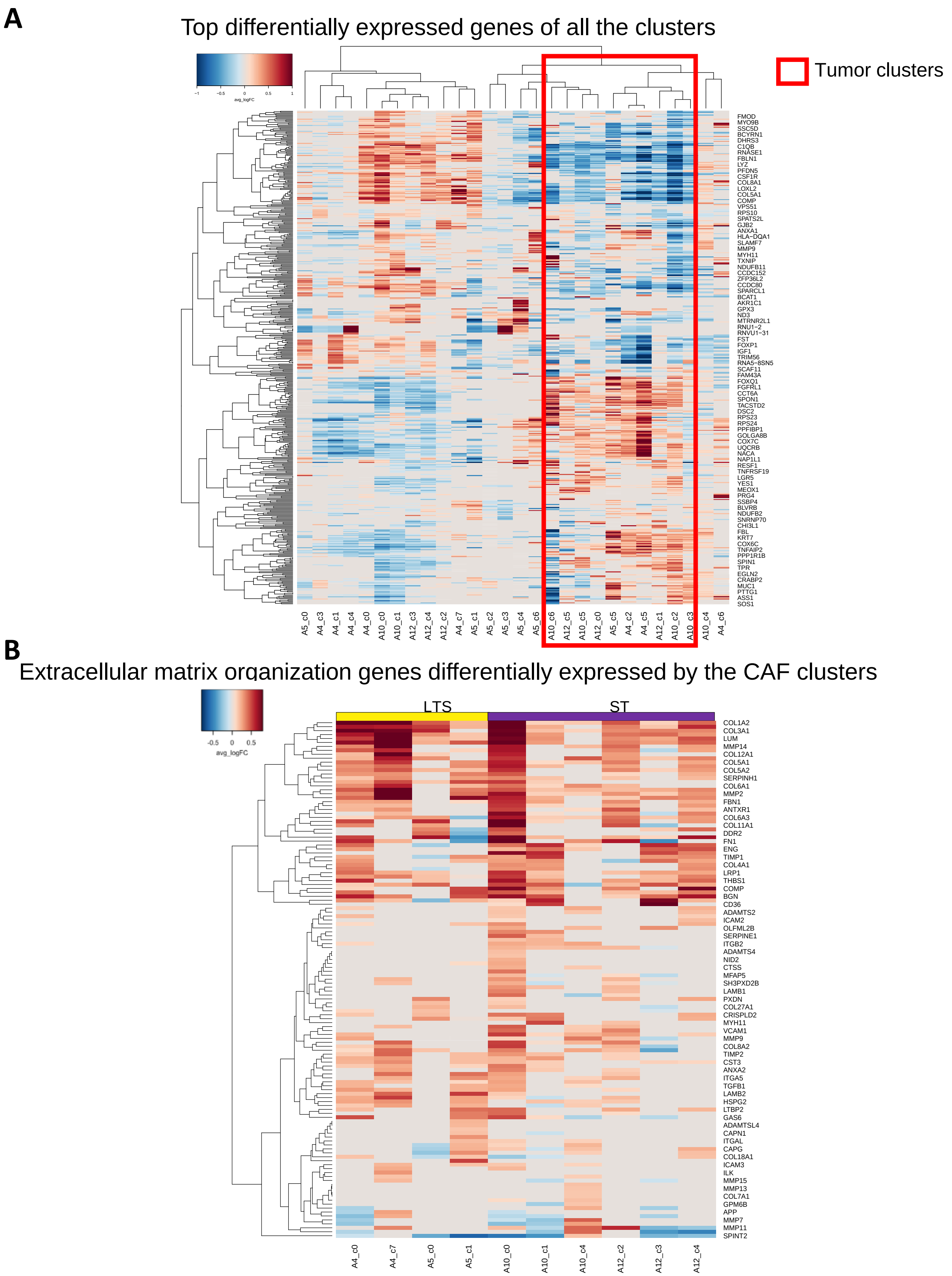


**Supplementary Fig. 1. Differentially expressed genes in spatial transcriptomics clusters. A.** The top 20 differentially expressed genes in each cluster (p < 0.01, Wilcoxon rank sum test; average log fold-change > 0.15). The average log fold-change was computed as the log fold-change of the average expression of a gene between a cluster and all other clusters of the sample. The red box indicates tumor clusters. **B.** Heatmap of the average log fold-change of the extracellular matrix organization genes (DAVID GO gene set) differentially expressed by the CAF clusters (p < 0.01, Wilcoxon rank sum test; average log fold-change > 0.15).

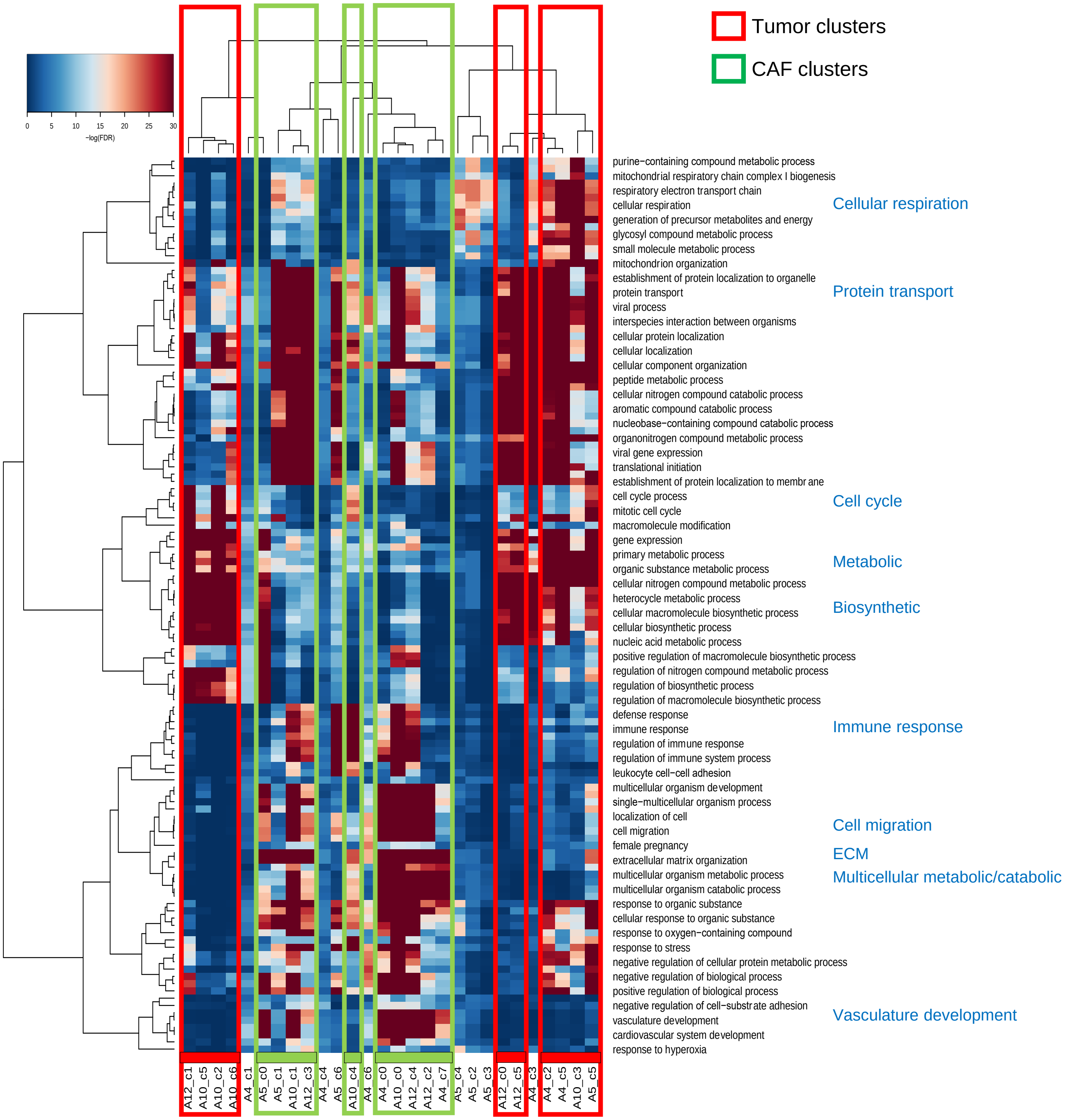


**Supplementary Fig. 2. Overrepresentation analysis of GO biological process gene sets.** Heatmap showing the –$\log_2$(FDR) of the hypergeometric test (n=20,000 background genes). The top 10 most enriched gene sets for each cluster were selected. Red boxes indicate the tumor clusters; green boxes indicate the CAF clusters.

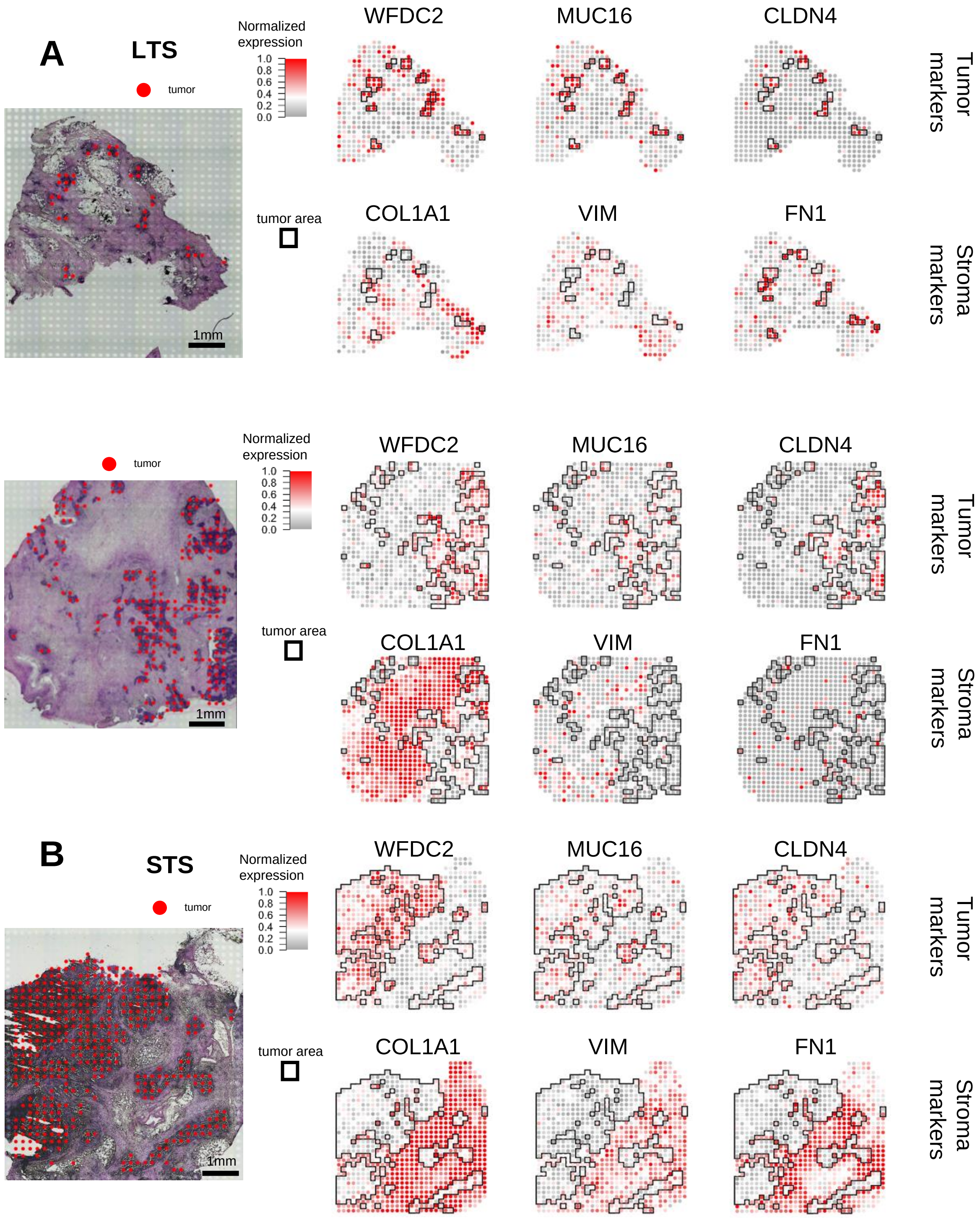


**Supplementary Fig. 3. Spatial expression of marker genes in spatial transcriptomics samples. A**. LTS sample. **B.** STS samples. Left, H&E staining. The tumor spots (red) identified by MIA are overlaid on the H&E images. Right, Normalized spatial expression of tumor, fibroblast, and immune markers. The black outlining indicates the tumor areas overlapping the red spots in the left panels.

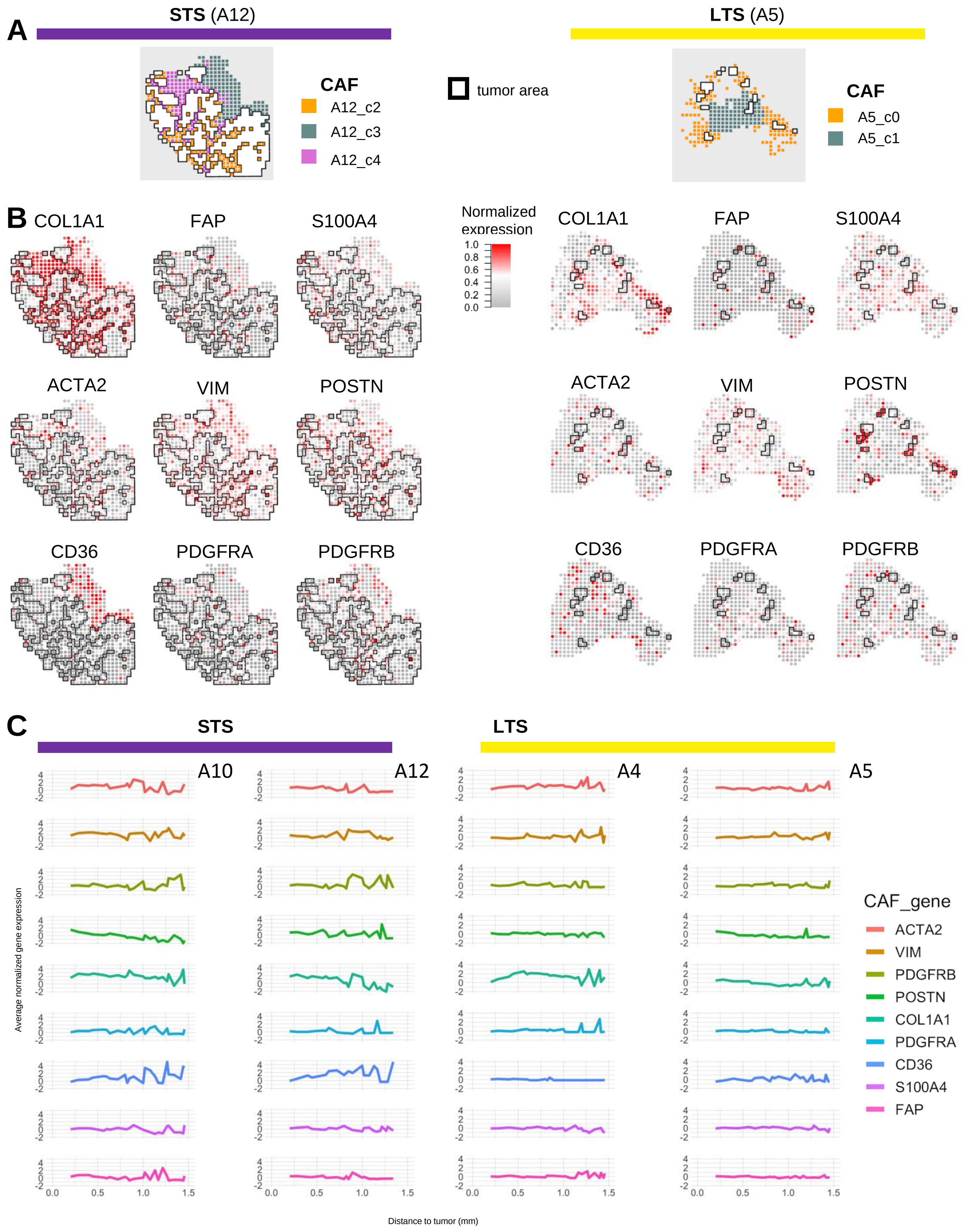


**Supplementary Fig. 4. Additional spatial maps for subtypes and distribution of CAF genes. A.** spatial mapping of CAF clusters in an STS sample (left) and an LTS sample (right)**. B.** spatial expression of selected CAF genes in an STS sample (left) and an LTS sample (right)**. C.** Average of normalized CAF gene expression as a function of the minimum distance to tumor in STS samples (left) and LTS samples (right).

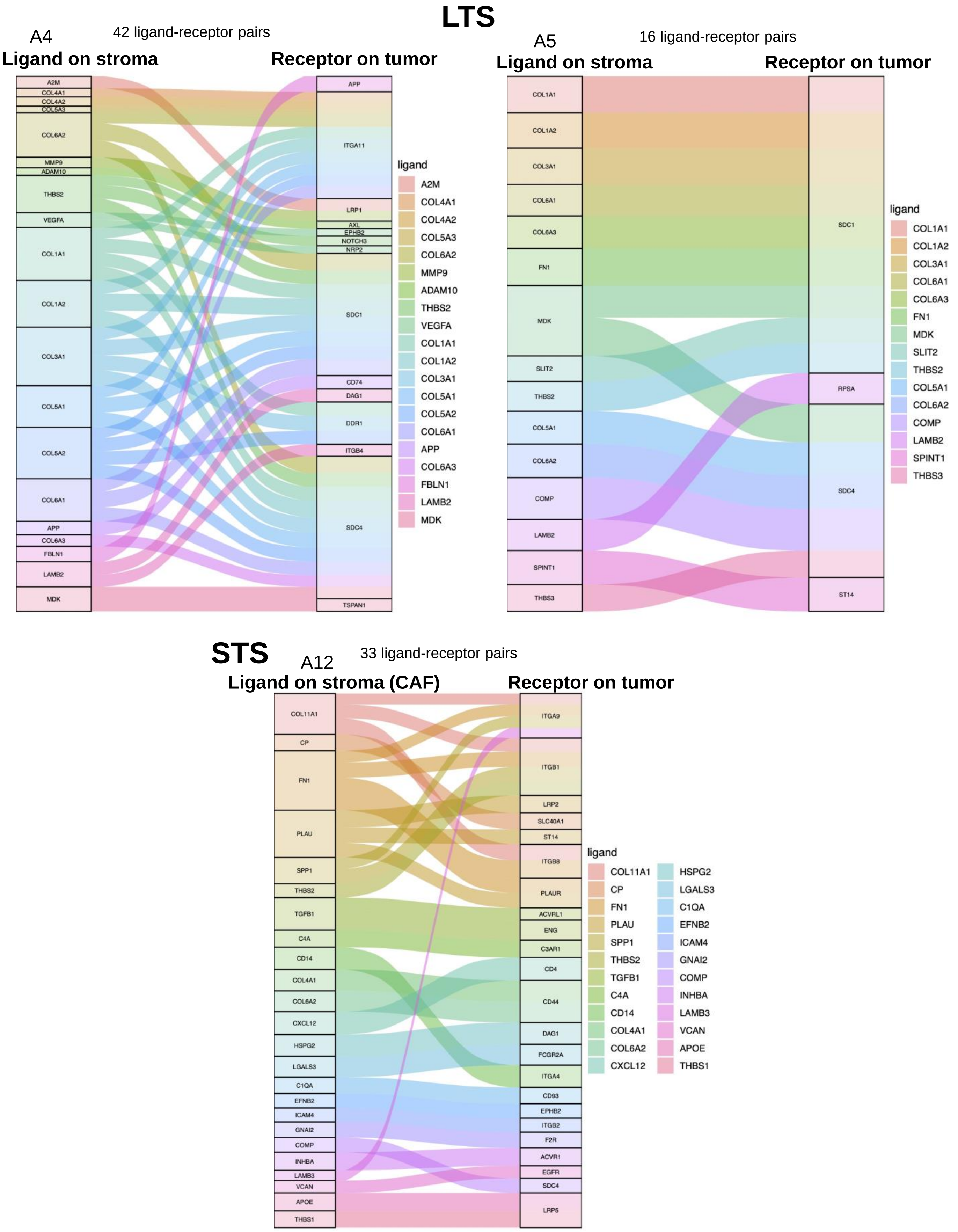


**Supplementary Fig. 5. Ligand-receptor interactions between stroma and tumor in additional samples.** Top, two LTS samples. Bottom, an STS sample. Ligand-receptor pairs identified at the stroma-tumor subregion interfaces and pooled together are shown.

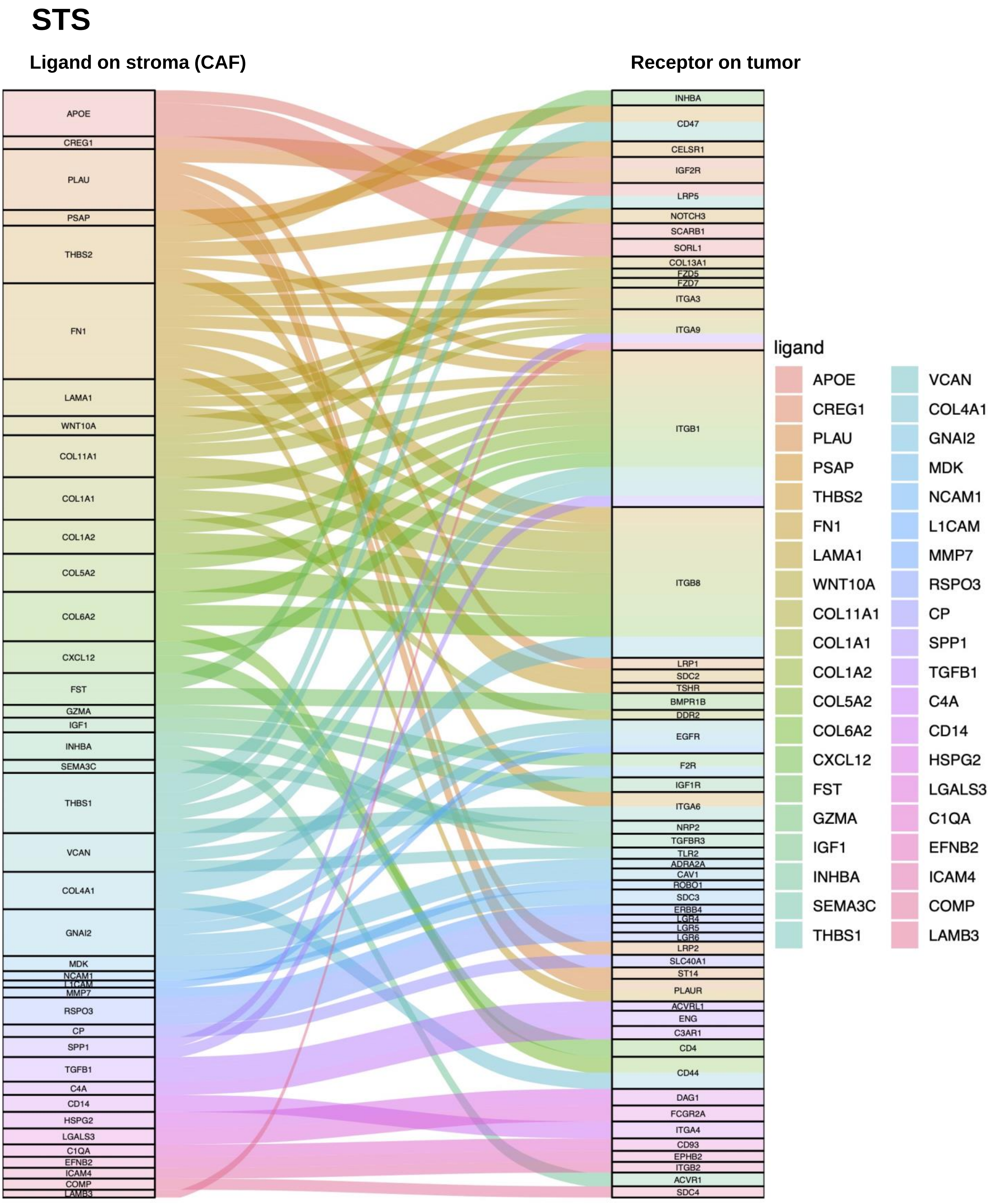


**Supplementary Fig. 6. Interactions between ligands on stroma and receptors on tumor in STS samples.** Eighty-four ligand-receptor pairs identified at all stroma-tumor subregion interfaces in two STS samples are shown.

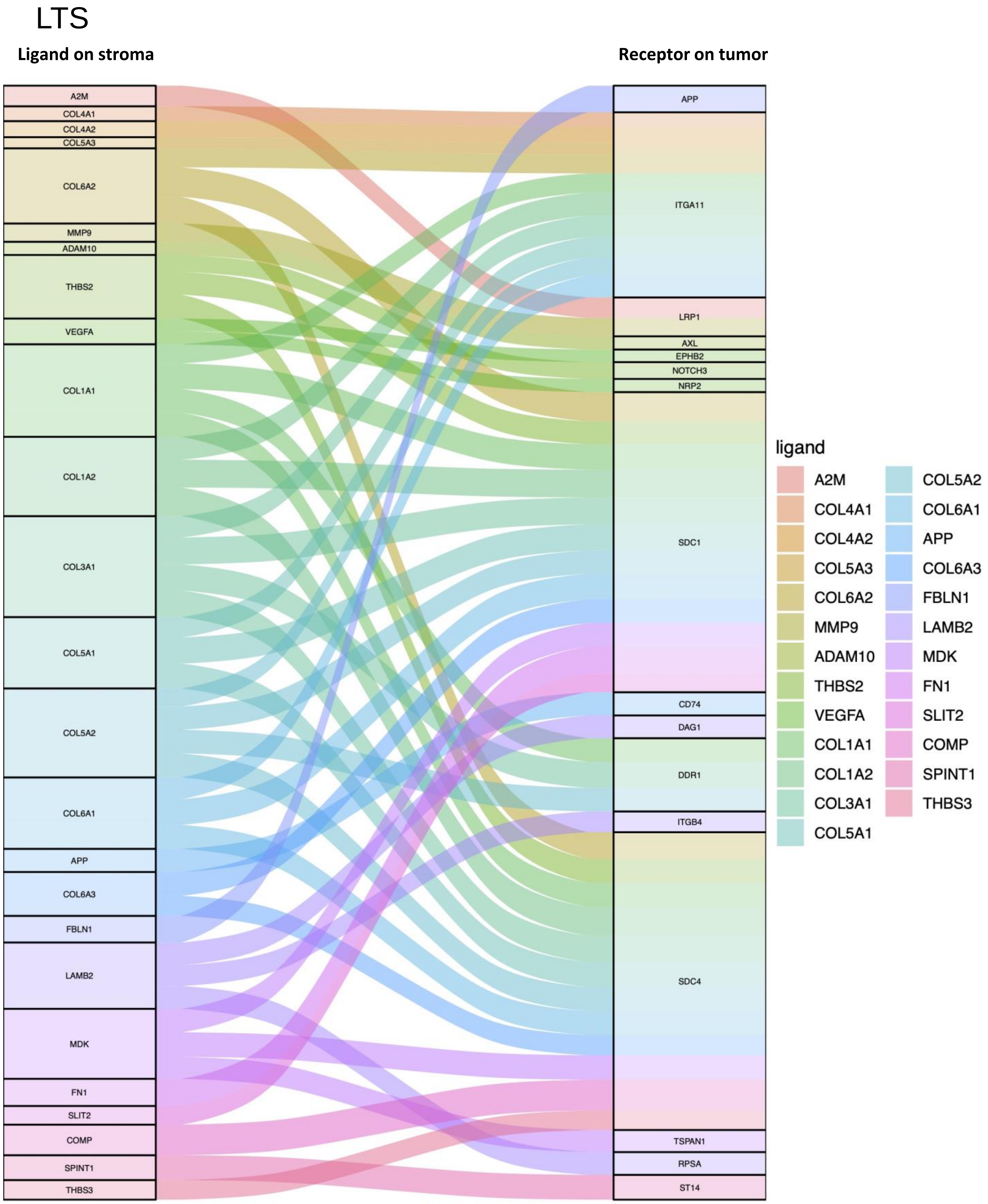


**Supplementary Fig. 7 Interactions between ligands on stroma and receptors on tumor in LTS samples.** Fifty ligand-receptor pairs identified at all stroma-tumor subregion interfaces in two LTS samples are shown.

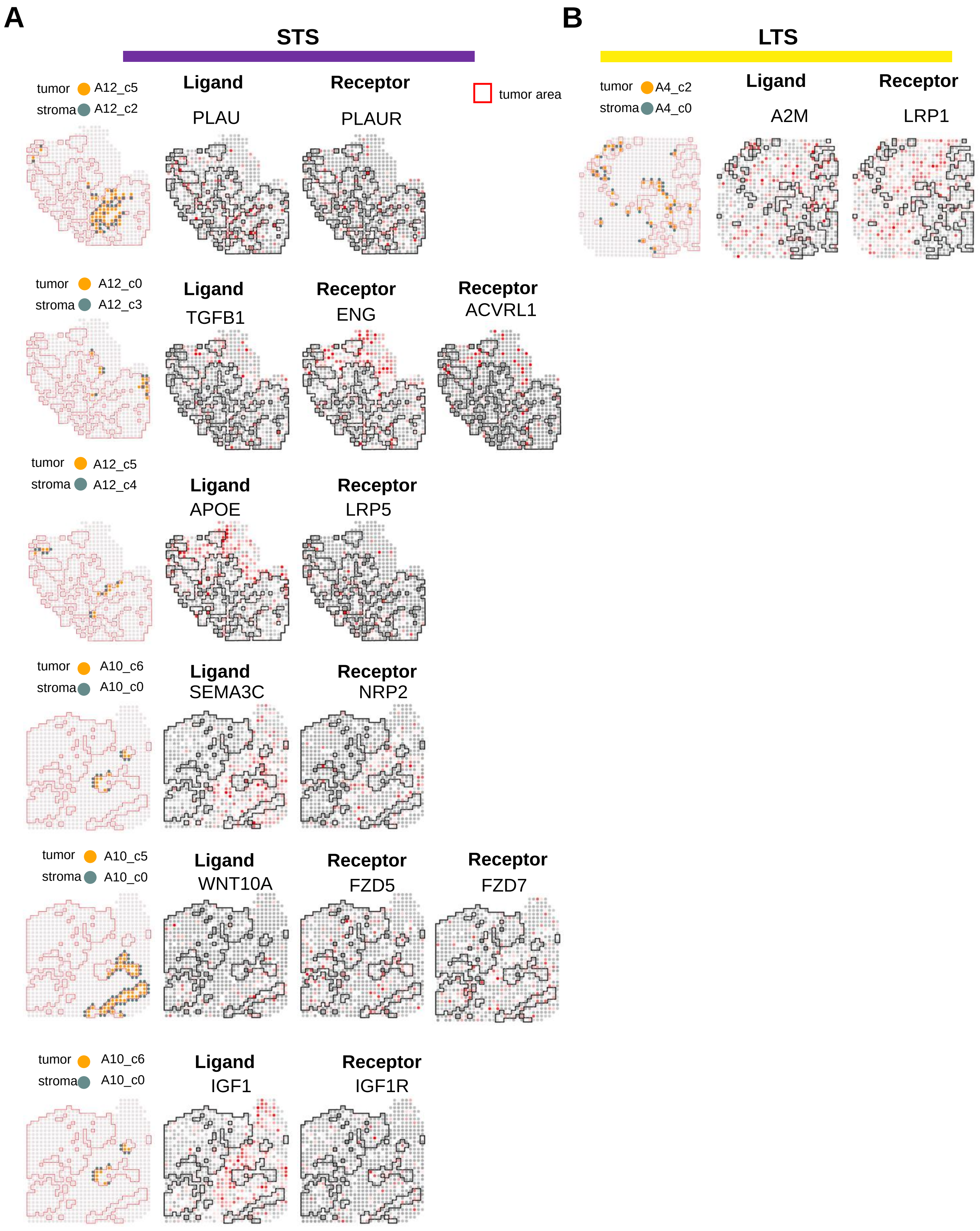


**Supplementary Fig. 8. Region-specific ligand-receptor interactions between stroma and tumor.** Representative nearest-neighbor interactions between tumor and stroma subregions in STS samples (**A**) and an LTS sample (**B**). Left, Localization of nearest-neighbor spots in the tumor subregion (orange) and stroma subregion (gray). Red outlining indicates the tumor area of the sample. Middle and Right, Spatial expression of ligand (middle)-receptor (right) pairs from stroma to tumor in the localized spots shown in the left panels. Black outlining indicates the tumor area of the sample.

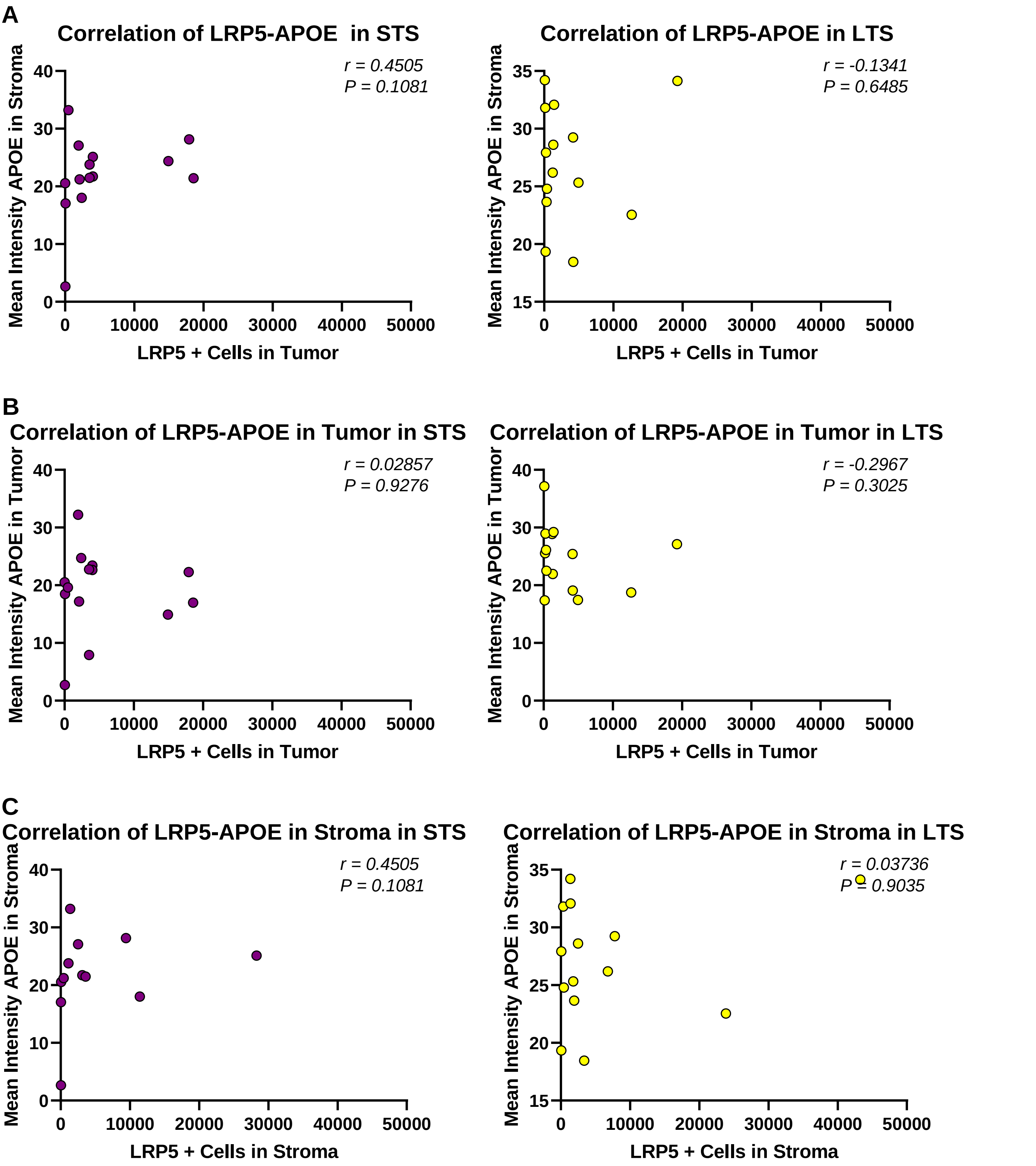


**Supplementary Fig. 9. Validation of ligand-receptor cross-talk in stroma and tumor. A.** Spearman correlation analysis of LRG5$^{+}$ cells in tumor and APOE expression intensity in stroma. **B.** Spearman correlation analysis of LRP5$^{+}$ cells and APOE expression intensity in tumor. **C.** Spearman correlation analysis of LRP5$^{+}$ cells and APOE expression intensity in stroma.
